# A-to-I RNA editing in SARS-COV-2: real or artifact?

**DOI:** 10.1101/2020.07.27.223172

**Authors:** Ernesto Picardi, Luigi Mansi, Graziano Pesole

## Abstract

ADAR1-mediated deamination of adenosines in long double stranded RNAs plays an important role in modulating the innate immune response. However, recent investigations based on metatranscriptomic samples of COVID-19 patients and SARS-COV-2 infected Vero cells have recovered contrasting findings. Using RNAseq data from time course experiments of infected human cell lines and transcriptome data from Vero cells and clinical samples, we prove that A-to-G changes observed in SARS-COV-2 genomes represent genuine RNA editing events, likely mediated by ADAR1. While the A-to-I editing rate is generally low, changes are distributed along the entire viral genome, are overrepresented in exonic regions and are, in the majority of cases, nonsynonymous. The impact of RNA editing on virus-host interactions could be relevant to identify potential targets for therapeutic interventions.

## Main Text

SARS-COV-2 is an enveloped virus with a positive sense, single-stranded RNA (ssRNA) genome of about 30 kb belonging to the genus *betacoronavirus* (Gorbalenya et al., 2020), sadly known for causing the pandemic by coronavirus disease 19 (COVID-19) (Poon and Peiris, 2020). Comparative genomics of thousands complete viral sequences of SARS-COV-2 from diverse geographic sites has revealed a biased subtitutional pattern in which the C-to-T change outnumbers all other substitutions (Chiara et al., 2020). The non-random occurrence of this mismatch strongly suggests that the SARS-COV-2 genome could undergo C-to-U RNA editing through APOBECs, as recently shown in metagenomic experiments from bronchoalveolar lavage fluids (BALF) of COVID-19 patients (Di Giorgio et al., 2020). On the other hand, there are contrasting evidences on the occurrence of A-to-I RNA editing, even though the A-to-G change appears the second most common mismatch type (Chiara et al., 2020). RNA editing by adenosine deamination is carried out by ADAR enzymes and is prominent in the human transcriptome in which it converts As in Is in long double-stranded RNAs (dsRNAs) formed by repeated elements in opposite orientation (mainly Alu sequences) (Eisenberg and Levanon, 2018). Human cells harbour three ADAR genes, ADAR (also known as ADAR1), ADARB1 (also known as ADAR2) and ADARB2 (also known as ADAR3) (Savva et al., 2012). ADAR1 and ADAR2 are expressed in almost all human tissues and are catalytically active (Picardi et al., 2015; Tan et al., 2017). While ADAR2 tends to edit As in coding protein sequences and only a few instances have been detected up to now (Pinto et al., 2014), ADAR1 extensively deaminates As in long dsRNAs and exists in two different isoforms, ADAR1p110, constitutively expressed, and ADAR1p150 mainly located in the cytoplasm and inducible by type I interferon (Savva et al., 2012).

Recently, it has been shown that A-to-G changes found in metagenomic sequences from BALFs of COVID-19 patients could be due to the activity of ADARs (Di Giorgio et al., 2020) but strong evidences of A-to-I RNA editing in the SARS-COV-2 genome have not been provided. Indeed, it is well known that ADAR1 tends to edit sites in clusters (hyper-editing) and exhibits a specific sequence context with G depletion one base upstream the edited site (Eisenberg and Levanon, 2018; Porath et al., 2014). These two important signatures were not detected in metagenomic sequences that, by their nature, prevent also the accurate quantification of ADARs as well as their RNA editing activity at transcriptomic level. Additionally, a concomitant study describing the transcriptome of SARS-COV-2 in infected Vero cells by using the nanopore direct RNA sequencing and the DNA nanoball sequencing, excluded ADAR mediated deamination for lack of A-to-G changes (Kim et al., 2020).

ADAR1 has a pivotal role in the modulation of the innate immune response, the first line defense against foreign viral nucleic acids (Lamers et al., 2019; Mannion et al., 2014). Through proteins called nucleic acids sensors, such as the endosomal Toll-like receptors (TLRs) and the cytoplasmic retinoic acid-inducible gene I (RIG-I) like receptors (RLRs), typical intermediates of virus replication, as dsRNA or ssRNA, can be recognized and induce the production of type I interferons (Koyama et al., 2008). In turn, type I interferons activate the expression of interferon-stimulated genes (ISGs), including ADAR1p150 and members of the APOBEC protein family (Borden and Williams, 2011). Once produced, ADAR1p150 can have antiviral effects by destabilizing dsRNAs through multiple A-to-G substitutions, an occurrence termed hyper-editing, or proviral effects suppressing the innate immune response by A-to-I RNA editing of long dsRNAs (Samuel, 2011, 2012). Consequently, exploring the origin of A-to-G changes occurring along the SARS-COV-2 genome could be quite relevant to better understand the host-virus relationships or the evolutionary dynamics of the viral genome and identify potential targets for therapeutic interventions.

Here we prove that A-to-G changes observed in the SARS-COV-2 genome are genuine RNA editing events likely mediated by ADAR1. By using an *ad hoc* computational workflow to mitigate the noise of sequencing errors, we were able to detect A-to-I editing in human and Vero infected cell lines as well as in several clinical samples.

We initially analyzed strand oriented paired end reads data from total RNA of Calu-3 human epithelial lung cancer cell line infected by SARS-COV-2 at a MOI of 0.3 (Emanuel et al., 2020). Total RNA was extracted at different time points post infection (4, 12 and 24 hours). Viral load was estimated as the fraction of reads mapping on the viral genome over the total number of reads per sample. Raw reads were cleaned to remove low quality regions and mapped on a comprehensive reference sequence including the whole human genome and the SARS-COV-2 genome (NC045512) by bwa. Unique and concordant SARS-COV-2 paired end reads were individually explored and filtered to detect reads carrying high quality A-to-G clusters (phred-score > 30). In all examined samples, we found a variable number of hyper-edited reads with a significant enrichment toward A-to-G and T-to-C clusters (on opposite strand) (Fig. 1A).

**Figure 1.**
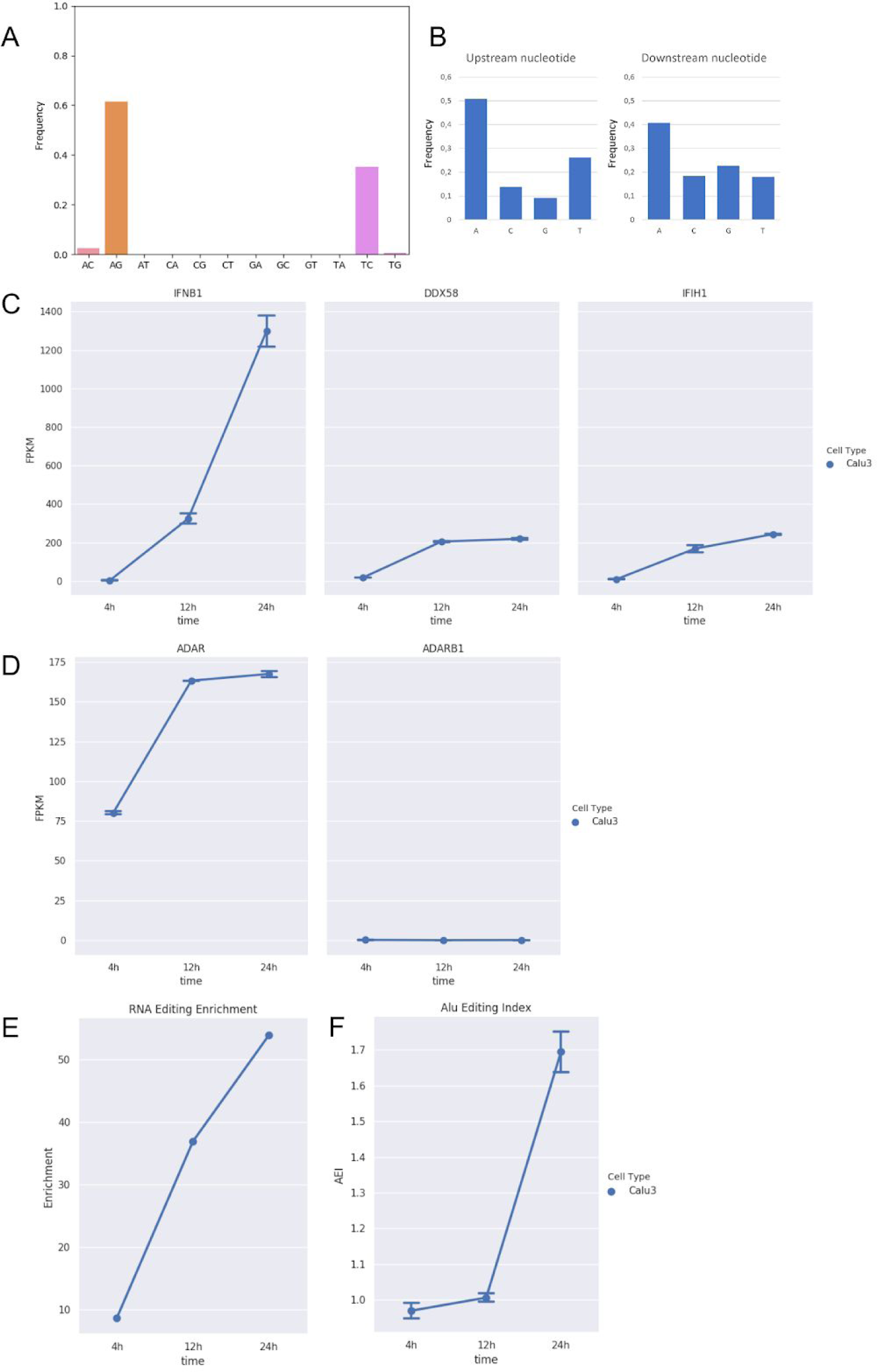
RNA editing and expression of key genes in RNAseq data from Calu-3 infected cells at three time points post-infection (4h, 12h and 24h). A) Distribution of hyper-editing identified in viral reads of infected Calu-3 cells. B) Nucleotide context one nucleotide upstream and downstream the detected hyper edited sites. C) Gene expression of type I interferon (*IFNB1*) and key sensor genes, *DDX58* (*RIG-I*) and *IFIH1* (*MDA5*) in infected Calu-3 cells. B) Expression of *ADAR* and *ADARB1* genes in infected Calu-3 cells. C) Enrichment of unique hyper editing positions in infected Calu-3 cells. D) Alu Editing Index (AEI) in infected Calu-3 cells.

While the majority of them were located on the sense strand, only a few A-to-G and T-to-C clusters were observed on the antisense strand, suggesting that A-to-I editing should mainly occur in long dsRNAs during the viral replication. On the whole, we detected 377 unique A-to-G events in 148 hyper-edited reads and their sequence context showed G depletion one base upstream and a slight G enrichment one base downstream of the editing sites, strengthening the evidence of ADAR1 mediated RNA editing (Fig. 1B).

During the infection and after the activation of type I interferon (IFNB1) (Fig. 1C) we observed an increased expression of ADAR1 (Fig. 1D), especially at 12 hour post infection (Pval < 0.05), and an enrichment in hyper-editing events (Fig. 1E). Such enrichment was marked at 24 hours post infection in which we registered a significant increment of the ADAR1 activity measured by the Alu editing index (AEI) (t-test Pval < 0.01) (Roth et al., 2019) (Fig. 1F). A finding indicating that ADAR1 and, in particular ADAR1p150, could be the main player of the observed A-to-I hyper editing.

Additionally, we analyzed strand oriented paired end reads data from total RNA of uninfected and SARS-COV-2 infected Vero cells in which no A-to-I editing events were detected (Kim et al., 2020). Vero cells derive from African green monkey fibroblasts that have lost the ability to produce interferon and are commonly used to grow interferon-sensitive viruses (Ammerman et al., 2008). By comparing RNAseq data of uninfected and infected Vero cells, we initially verified the absence of the type I interferon response (IFNB1) to the viral infection and the expression of ADAR1 (Fig. 2AB).

**Figure 2.**
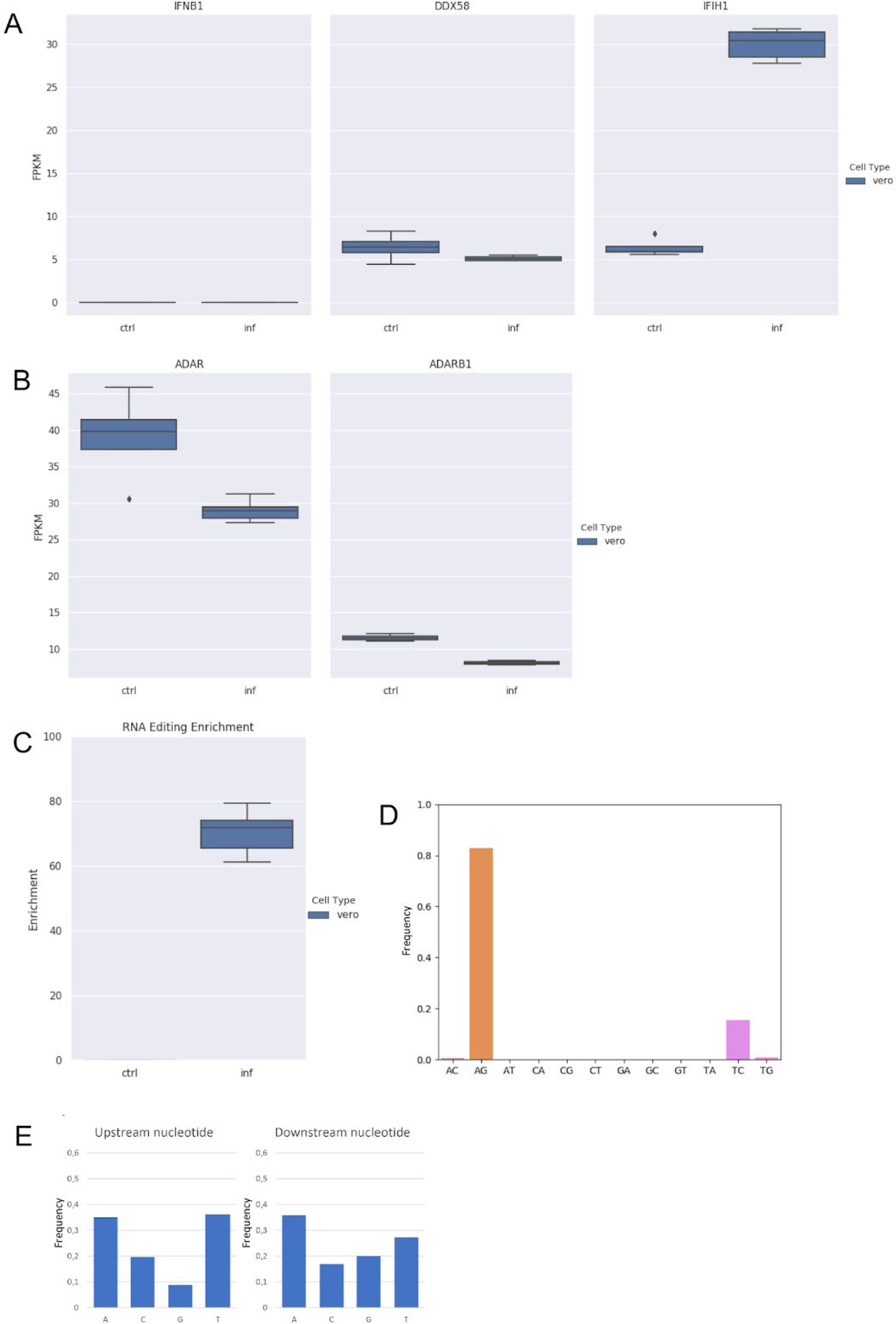
RNA editing and expression of key genes in RNAseq data from infected (inf) and uninfected (ctrl) Vero cells. A) Gene expression of type I interferon (*IFNB1*) and key sensor genes, *DDX58* (*RIG-I*) and *IFIH1* (*MDA5*) in Vero cells. B) Expression of *ADAR* and *ADARB1* genes in Vero cells. C) Enrichment of unique hyper editing positions in Vero cells. D) Distribution of hyper-editing identified in viral reads of Vero cells. E) Nucleotide context one nucleotide upstream and downstream the detected hyper edited sites.

Next, by applying the above described computational strategy, we found 1207 hyper edited reads (∼201 per sample) enriched in A-to-G and T-to-C clusters (98% of all hyper edited reads) (Fig. 2CD), even though ADAR1 appeared downregulated upon the infection (Fig. 2B). A-to-I editing was enriched at the same level in all replicates of infected Vero cells (Fig. 2C) and the sequence context showed G depletion one base upstream of the editing sites (Fig. 2E), indicating that also in Vero cells the SARS-COV-2 genome undergoes A-to-I RNA editing.

We observed a strong correlation (Pearson R 0.97 - Pval << 0.01) between the number of hyper edited reads and the number of viral reads, justifying the highest number of hyper edited reads in Vero cells, despite the lack of type I interferons. Indeed, an average of 10M of viral reads were used in Calu-3 against an average of 54M in Vero cells.

RNA editing and the activity of ADAR1 are tissue and cell type dependent as well as the type I interferon response to the viral infection (Picardi et al., 2015, 2017). To investigate A-to-I editing in different SARS-COV-2 infected cell lines, we analyzed stranded PolyA+ single end RNAseq data from Calu-3, Caco-2 and H1299 human cell lines infected by SARS-COV-2 at a MOI of 0.3 and generated at three time points (4, 12 and 24 hours) post infection (Emanuel et al., 2020). The viral infection activated the type I interferon response in Calu-3 cells only and, consequently, ADAR1 did not appear deeply up regulated in Caco-2 and H1299 cells as also attested by the AEI index measured at all time points (Supp. Fig. 1). We found A-to-G and T-to-C hyper edited reads only in Calu-3 and Caco-2 cells but the total number of edited reads was quite low as a result of the PolyA+ sequencing strategy in which mature viral transcripts rather than full genomic RNAs are captured. Additionally, viral reads from PolyA+ data were about 4 orders of magnitude less abundant than total RNAseq data.

In parallel, we profiled RNA editing at single nucleotide resolution using the strategy described by Di Giorgio et al. (Di Giorgio et al., 2020) but implementing more stringent filters. We used only concordant paired reads whose alignments were confirmed by two independent mappers (bwa and gsnap). Single nucleotide variants detected by REDItools (Picardi and Pesole, 2013) were called at an allelic fraction two times higher than the error rate estimated by the overlap of read pairs. Strand biases were corrected employing the strand oriented protocol for sequencing. In Calu-3 total RNA data, we found 756 putative A-to-I events, increasing from 4 hours to 24 hours post infection and accounting for about 35% of all nucleotide variants. Interestingly, about 42% of all base changes were C-to-T substitutions most likely due to the APOBECs activity. In infected Vero cells, we detected 741 A-to-I candidates but we did not observe an enrichment in A-to-G and T-to-C events. As in Calu-3 cells, C-to-T changes outnumbered the majority of inferred substitutions even though the G-to-A change emerged as prominent. In PolyA+ data, instead, only a tiny number of nucleotide variants was detected and again C-to-T appeared the most representative substitution.

As already shown in (Di Giorgio et al., 2020), A-to-I candidates as well as C-to-U candidates displayed very low editing levels, less than 1% in more than 99% of positions. Also Alu repetitive elements in the human transcriptome tend to be edited at extents lower than 1% (Picardi et al., 2016; Roth et al., 2019) and this strengthens the idea that ADAR1 should be the main player of the SARS-COV-2 adenosine deamination. However, differently from sites in hyper edited reads, events detected by this strategy should be regarded with high care. While in the human transcriptome A-to-G changes due to RNA editing can be distinguished from SNPs by means of whole genome (WGS) and/or whole exome (WES) sequencing data (Diroma et al., 2019), in the SARS-COV-2 RNA genome this distinction cannot be done. Although RNA editing modulation observed in time-course experiments is a remarkable evidence, genuine RNA editing substitutions cannot be easily discerned from nucleotide variants due to sequencing or polymerase errors.

Finally, we re-analysed metagenomic samples already used in (Di Giorgio et al., 2020) but limiting our workflow to samples in which only paired end reads were available (Supp. Table 1). In BALF samples from bioproject PRJNA605907 (Shen et al., 2020) we found hyper editing enrichment only in experiments with a high number of viral reads (>4.5M). Such samples showed a deep coverage depth of the viral genome (also > 7000x) and more than 75% of detected substitutions were A-to-I candidates. In the two metagenomic BALF samples from bioproject PRJNA601736, we were unable to identify hyper edited reads and only a few putative RNA editing sites were detected at single nucleotide level. In these samples, however, the coverage depth of the viral genome was relatively low (103x and 430x) as well as the number of viral reads (∼65000 on average).

We also analyzed RNAseq samples from bioproject PRJNA616446 including reads from BALFs and throat swabs. We detected a few hyper edited reads only in BALF samples and the distribution of nucleotide variants was in line with previous observations from metagenomic samples. Viral genomes from throat swabs were supported by a low number of reads. We also tried to detect RNA editing in RNAseq experiments from post mortem human donors of the bioproject PRJNA631753 but the number of viral reads per sample was too small to infer high quality A-to-I events. Together, our results from infected cell lines and clinical samples show clear A-to-I editing signatures in the SARS-COV-2 genome, even though its reliable profiling is strictly dependent on the sequencing strategy and number of viral reads (in turn related to the viral load). In all cases, the impact of ADAR mediated RNA editing on the SARS-COV-2 genome, in terms of A-to-I events or hyper edited reads as well as editing frequency, is generally low and most likely due to the following factors: 1) absence of very long dsRNAs along the viral genome or subgenomic regions that could be firmly bound by ADAR1 (Rangan et al., 2020); 2) dsRNAs from intermediates of viral replication, that are expected to be the optimal targets of ADARs, are poorly represented and the antisense strand is less than 1% as abundant as the sense counterpart (estimated by stranded RNAseq data); 3) viral RNA synthesis is associated to double-membrane vesicles preventing the action of cytoplasmic enzymes (Snijder et al., 2020); 4) the viral RNA-dependent RNA polymerase has proofreading activity that could mitigate the effect of deamination (Romano et al., 2020); 5) SARS-COV-2 seems to have mechanisms to evade and suppress the interferon response, leading to low induction and expression of antiviral genes (including ADARs and APOBECs) (Park and Iwasaki, 2020).

Taking all unique A-to-I editing events detected in hyper edited reads of all analysed cell lines, representing the high quality fraction of edited SARS-COV-2 sites, we discovered that they are distributed along the entire viral genome with an apparent preference towards the 3’ end region (Supp. Fig. 2), a finding suggesting that genomic dsRNAs are indeed the main SARS-COV-2 double strand structures targeted by ADAR1.

The 96% of sites from hyper edited reads reside in exonic viral regions and comprise 64% of nonsynonymous events that could have a strong functional impact on the SARS-COV-2 pathogenicity.

We cannot establish how the virus escapes the antiviral action of RNA editing but several events are fixed and maintained. Although RNA editing occurs at low extent, it could be one of the most relevant mechanisms governing the dynamics of viral evolution. In addition, edited variants could significantly influence the virulence, pathogenicity and host response. Since the virus tends to evade the RNA editing action, it means that it could have a strong impact on its survival. On the other hand, RNA editing is emerging as a promising therapeutic alternative for different human genetic disorders (Katrekar et al., 2019; Reardon, 2020) and, thus, it could have an important role in the antiviral fight against the SARS-COV-2 and/or other RNA viruses.

## Acknowledgments

We thank Thomas Wu for fixing and enabling the gsnap transcriptome mapping option for SARS-COV-2. We also thank ELIXIR Italy and the ReCaS calculus centre at the University of Bari for providing the computing and bioinformatics facilities.

## Author Contributions

E.P. conceptualized the study, wrote main scripts and drafted the manuscript. L.M. performed bioinformatics analyses. G.P. supervised the work and edited the final version of the manuscript.

## STAR★Methods

### Resource Availability

#### Lead Contact

Further information and requests should be directed to and will be fulfilled by the Lead Contact.

#### Materials Availability

This study did not generate new unique reagents.

#### Data and Code Availability

The raw data are available at SRA under the following BioProject accessions: PRJNA625518, PRJNA616446, PRJNA601736, PRJNA605907 and PRJNA631753. The raw data from infected Vero cells are available at the Open Science Framework (OSF) with accession number https://doi.org/10.17605/OSF.IO/8F6N9, while data from uninfected Vero cells are available at SRA with accessions: DRR018832, DRR018833, DRR018834 and DRR018835.

Scripts used to detect RNA editing events are available at the following GitHub link https://github.com/BioinfoUNIBA/sars-cov-2-editing.

### Method Details

#### Filtering of RNAseq raw data

Raw reads were cleaned using FASTP (Chen S, 2018) and taking into account the read length. For reads longer than 76 bases, we trimmed 10 nucleotide upstream and downstream, and removed reads with more than 20% of unqualified bases (-q 25 -u 20 -l 50 –x --cut_tail --cut_tail_mean_quality 25 --trim_front1 0 --trim_tail1 0). The trimming was disabled for reads shorter than 76 bases. The mean quality per base was fixed at a phred-score of 25. Reads shorter than 50 bases were removed.

#### Alignment of RNAseq reads

Cleaned reads were aligned onto a comprehensive reference sequence including the whole human genome (hg19 assembly from UCSC) and the SARS-COV-2 genome (NC045512.2 from NCBI) by bwa (Li and Durbin, 2009) using default parameters. Unique and concordant reads mapping on the SARS-COV-2 genome were extracted by sambamba (Tarasov et al., 2015) and converted in BAM format by SAMtools (Li et al., 2009). Viral reads were also aligned onto the NC045512.2 assembly by GSNAP (Wu and Nacu, 2010) employing the transcriptome-guided strategy. SARS-COV-2 transcript annotations were obtained from UCSC. The strand orientation per each sample was inferred by the infer_experiment.py script from the RSeQC package (Wang et al., 2012). Additionally, human reads were also aligned onto the human reference genome by STAR (Dobin et al., 2013) and proving known GENCODE (v31lift37) annotations.

In Vero cells, the human genome was replaced by the *Chlorocebus sabaeus* genome (chlSab2 assembly) from UCSC. Green monkey annotations were downloaded also from UCSC.

#### Detection of hyper edited reads

The detection of hyper edited reads was performed using our custom SubstitutionsPerSequence.py script. It takes as input viral reads aligned by bwa (Li and Durbin, 2009) in BAM format and filters out reads with a mapping quality score lower than 30, not properly mapped, flagged as secondary alignments, carrying insertions or deletions and showing more than 2 substitutions of different type. Reads with mismatches of an unique type are further filtered in a similar way as described in (Porath et al., 2014). Dense clusters of high-quality (Phred ≥30) A-to-G (or T-to-C) mismatches are detected retaining reads in which the number of A-to-G changes was at least 5% of the read length and discarding reads having too dense A-to-G mismatch clusters (length <10% of the read length) or clusters contained within either the first or last 20% of the read or clusters with a particularly large percentage (>60%) of a single nucleotide. Reads in which the aligned region was less than 80% of the read length are also removed.

#### Detection of RNA editing at single nucleotide level

We performed an initial variant calling by REDItools (version 2) (Picardi and Pesole, 2013) and same parameters used also in (Di Giorgio et al., 2020) (-os 4 -q 30 -bq 30 -l 0). Strand orientation was taken into account in samples in which libraries were prepared using strand oriented kits. To remove the noise due to sequencing errors, we used only concordant reads whose alignments were confirmed by two independent aligners, bwa (Li and Durbin, 2009) and gsnap (Wu and Nacu, 2010). In addition, we excluded discordant base variants at overlapping regions of read pairs. Low quality reads and reads not properly mapped or flagged as secondary alignment or carrying indels were removed as well. We excluded a certain number of positions in the first of last regions of reads depending on the read length (5 upstream and 6 downstream for reads < 100 bases, 15 upstream and downstream for reads > 150 bases). All sites in which the variant nucleotide was not supported by at least 4 reads were removed. In strand oriented experiments, the variant calling was corrected accordingly. In non strand oriented experiments, instead, the fisher exact test was used to check strand biases.

In the variant calling we excluded also positions in single repeats and known viral variants from UCSC. The entire procedure of noise correction was implemented in the corr.py script.

After the noise correction, RNA editing candidates were called at a minimal allele frequency equal to two times the error rate estimated by the overlaps of read pairs.

#### Gene expression in cell lines

Read counts per known gene were carried out using featureCounts (Liao et al., 2014) and GENCODE (v31lift37) annotations. Differential expression in time course experiments was done by DESeq2 (Love et al., 2014) while count normalization in FPKM for figures was performed by a custom script.

#### Alu editing index

The Alu editing index (AEI) proding the ADAR activity at transcriptome level was calculated using the pipeline described in (Roth et al., 2019). Differential AEI was assayed by the t-test at a significant level of 0.05.

#### Quantification of sense and antisense viral strands

The quantification of sense and antisense viral strands was performed in strand oriented datasets only and using featureCounts (Liao et al., 2014) providing as annotations the list of known viral non overlapping coding regions from UCSC. The percentage of the antisense viral strand was calculated as the fraction of reads mapping on coding sequences projected on the antisense strand over the total number of reads mapping on non overlapping coding sequences.

#### RNA editing enrichment

RNA editing enrichment was calculated taking into account only unique A-to-I events detected in hyper edited reads and according to the definition proposed by (Porath et al., 2014) in which the editing enrichment is equal to the number of unique A-to-I events in each experiment divided by the expected number. Such expected number was computed by multiplying the total number of A-to-I events (over all experiments) by the ratio of the number of mapped reads in the experiment to the number of mapped reads in all experiments, normalized by the viral load.

#### Annotation of A-to-I editing events

RNA editing events were annotated using ANNOVAR (Wang et al., 2010) providing the list of known SARS-COV-2 transcripts from UCSC.

## Supplemental Information

**Supplementary Figure 1.**
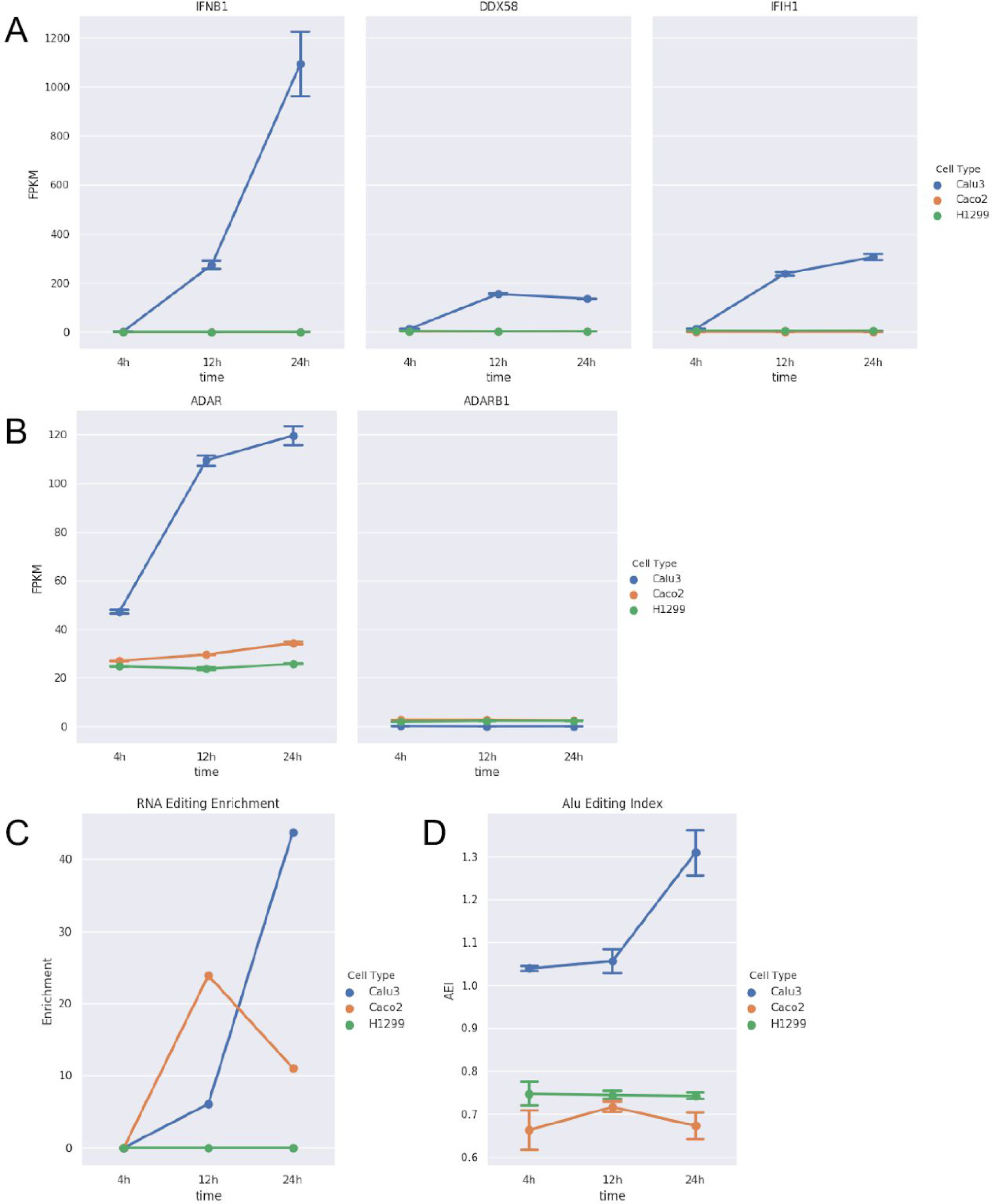
RNA editing and expression of key genes in PolyA+ RNAseq data from Calu-3, Caco-2 and H1299 infected cells at three time points post-infection (4h, 12h and 24h). A) Gene expression of type I interferon (*IFNB1*) and key sensor genes, *DDX58* (*RIG-I*) and *IFIH1* (*MDA5*). B) Expression of *ADAR* and *ADARB1* genes. C) Enrichment of unique hyper editing positions across infected cell lines. D) Alu Editing Index (AEI) in all cell lines.

**Supplementary Figure 2.**
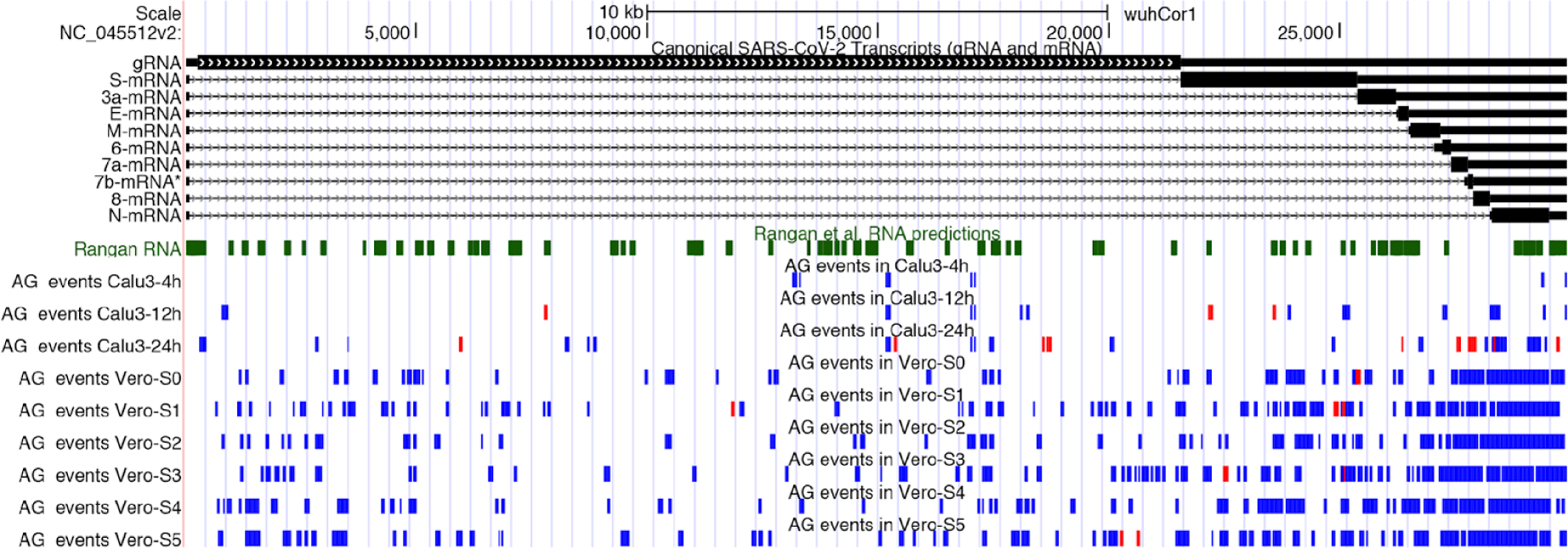
Detected SARS-COV-2 RNA editing events in their genomic context.

**Supplementary Table 1.** Statistics and RNA editing detected in clinical samples.

